# Varying richness need not imply non-random species co-occurrence: implications for specifying null models

**DOI:** 10.1101/2020.11.04.367839

**Authors:** Tom M. Fayle

## Abstract

**Background:** Non-random species co-occurrence is of fundamental interest to ecologists. One approach to analysing non-random patterns is null modelling. This involves calculation of a metric for the observed dataset, and comparison to a distribution obtained by repeatedly randomising the data. Choice of randomisation algorithm, specifically whether null model species richness is fixed at that of the observed dataset, is likely to affect model results. This is particularly important in cases when there is high variation in species richness between sampling units in the observed data.

**Methods:** Here I demonstrate the effects of accounting for variation in species richness. I use the C-score, a metric measuring species segregation as “checkerboard units”, applied to 289 datasets. First, I run null models in which sites are equally likely to be occupied (fixed-equiprobable algorithm). I do this both for the original datasets, and for the same datasets where occurrences are randomised with the species richness distribution fixed (pre-randomised datasets). Second, I run null models that fix site species richness to that observed (fixed-fixed algorithm).

**Results:** For real datasets, using the fixed-equiprobable algorithm (sites are equally likely to be colonised), C-score standardised effect size (SES) was positively related to variability in species richness between sites within a dataset. This effect was also found for pre-randomised datasets, indicating that variability in species richness can be exclusively responsible for detection of non-random species co-occurrence. When using the fixed-fixed algorithm (richness is constrained to that of real sites), there was no relationship between SES and variability in species richness. There was also a reverse in the effect direction, with 94% of significant tests indicating a lower C-score than expected for the fixed-equiprobable algorithm, but 98% of significant tests indicating a *higher* C-score than expected for the fixed-fixed algorithm.

**Discussion:** I speculate that when variation in species richness is high, fewer checkerboard units are possible, regardless of segregation between species. Therefore, use of fixed-equiprobable algorithms in situations where real species richness is highly variable between sites within a dataset will yield significant results, even if species co-occur randomly within the constraints of the species richness distribution. Consequently, use of such tests makes the a priori assumption that high within-dataset variation in species richness indicates non-random species co-occurrence. I recommend using algorithms that explicitly take into account species richness distributions when one wants to eliminate the effect of richness variation in terms of producing significant but spurious positive co-occurrence results. Alternatively, non-null mechanistic models can be created, in which hypothesised species assembly processes must be explicitly stated and tested.

## Introduction

Species commonly co-occur non-randomly with respect to each other. Determining the particular deterministic patterns to which species conform lies at the heart of ecology. Non-random co-occurrence patterns can be caused by a range of interactions between species, including competition, predation, parasitism, facilitation, and mutualism (Bell et al. 2010; Holdaway & Sparrow 2006). Drivers other than direct interactions can also play a role, including similar or differing environmental preferences leading to positive or negative co-occurrence respectively, and dispersal limitation over different scales (Holt et al. 2013; Hubbell 2001). Often data for testing such patterns take the form of a presence/absence species by sites matrix. One approach for analysing such matrices is to use null models of species co-occurrence (Gotelli 2000). This suite of methods involves the calculation of a metric measuring the way in which species co-occur. However, solely from such a metric it is not possible to tell whether the structure of the matrix differs from that which would be expected if species co-occurred independently. For example, even in randomly structured matrices it is likely that some pairs of species will not co-occur (Connor & Simberloff 1979). To overcome this problem, observed species occurrences are randomised to break any associations between the species. This is done repeatedly, and the metric calculated for each of these randomised matrices, allowing estimation of the distribution of the metric expected under the null hypothesis that species co-occur independently of each other (hence the term “null modelling”). This then allows the calculation of a p-value, that is, the probability that the observed metric, or one more extreme, would be observed under the null hypothesis of random species co-occurrence.

A critical step in this null modelling process is defining what is “random” in a given situation, and hence choosing exactly how to create randomised matrices from the observed data (Gotelli & Ulrich 2012). However, this aspect of the field remains contentious (e.g. Fayle & Manica 2011; Gotelli & Ulrich 2011). Specifically, the modeller needs to select which aspects of the real dataset to maintain in the null model.

Algorithms range from those that do not constrain either species richness or species occurrences across sites, keeping constant only the total number of species occurrences across the entire matrix, to more restrictive methods that constrain both richness and species occurrence distributions to be exactly the same as those from the observed dataset (Connor & Simberloff 1979; Gotelli 2000; Miklós & Podani 2004). Most analyses constrain the null occurrences of species to be the same as those observed in the real population (Fayle & Manica 2010), thus assuming that the occurrences of species in the observed sample correlate with the occurrences of species in the unobserved species pool. However, deciding the degree to which null species richness should reflect that of the real community is more problematic (Wilson 1995). Either null species richness across sites can be constrained to exactly match those observed (or some probabilistic version of this), or all sites can be defined to be equally likely to be occupied by a particular species, although with the richness distribution being able to vary to some degree (Gotelli 2000).

Constraining species richness values has the disadvantage that differences in species richness might have been caused by non-random co-occurrence between species. For example, presence of a keystone mutualist might lead to highly variable species richness, dependent on the presence of that species. Since this is the very mechanism that is under study it should therefore be excluded from the null model, potentially making any test that includes such constraints overly conservative (i.e. risking type 2 errors). This has been called the “Narcissus effect” (Colwell & Winkler 1984), since the signal of the processes under study is already “reflected” in the null model, and hence is not detected.

However, allowing unconstrained occupation of sites means that factors causing non-random differences in species richness between sites unrelated to species co-occurrence patterns are also excluded from the null model (the “Jack Horner effect”, Wilson 1995). Thus, it is not clear whether significant results of analyses using this less conservative method arise specifically from non-random species co-occurrence patterns, or merely from the real species richness distribution being more variable across sites within a dataset than is accounted for in the null model (in which case significant results could be type 1 errors). An example of this is the need to account for differences in island size when assessing co-occurrence between island dwelling species, since larger islands will have more species, because they have more living space and also present a larger target for incoming colonists (Connor & Simberloff 1979). Even if all islands represented identical habitats in terms of abilities of different species to survive there, it would still be necessary to account for differences in island size in a null model of species co-occurrence.

This problem was discussed during earlier development of null modelling of species co-occurrence (Colwell & Winkler 1984; Wilson 1995). However, sometimes a range of different methods for generating null matrices is used, and any non-random patterns that emerge are discussed without reference to null model constraints (e.g. De los Ríos et al. 2008). It is even possible to get opposite patterns (segregation vs aggregation) by using different randomisation algorithms on the same dataset (Hétérier et al. 2008; Rooney 2008), which is clearly a concern. The differences in results are expected to be more severe the greater the variation in species richness between sites within a dataset, since both randomisation methods should give similar results for datasets with low variation in species richness, because the null models specified will be similar. Note that co-occurrence metrics (even when standardised using null models) are correlated with the richness of the entire assemblage across all sites (Ulrich et al. 2017; Ulrich et al. 2018). Hence it is recommended that analyses including datasets with varying total species richness (and number of sites), account for this, at a minimum by using these measures of matrix size as a covariate during modelling (Ulrich et al. 2018). However, the effects of variation in species richness between sites *within* an assemblage remain unexplored. The purpose of this article is to demonstrate how variation in site species richness (as opposed to total assemblage richness across all sites) impacts the results of null models of species co-occurrence, with parallel analyses using randomisation algorithms that either constrain or do not constrain site species richness. Note that here I use “unconstrained species richness” to mean that null model species richness is not dictated to be precisely that of the observed dataset; I do not mean that species richness is completely free to vary.

I first quantify the degree to which within-dataset variability in species richness correlates with the detection of significant patterns of species segregation in 289 real datasets when that richness variability is not taken into account in the null model. To assess the degree to which this pattern might relate solely to variability in species richness itself, I then generate a series of datasets that match the real ones in terms of species richness and species occurrence distributions, but for which species occurrences are already randomised, and repeat this analysis. If detected patterns of species segregation relate only to variability in species richness (and not to non-random patterns of species co-occurrence), then we would expect the results from these two analyses to be similar. Finally, I repeat both analyses using a randomisation algorithm that constrains null site species richness to match the species richness distribution of the real datasets. These models should allow detection of species segregation or aggregation within the observed pattern of species richness.

## Materials and methods

### Does variability in species richness between sites within a dataset correlate with detection of deterministic species co-occurrence patterns?

First, I tested the degree to which variability in species richness affects the probability of finding significant patterns of co-occurrence between species in real datasets when this observed variability is not included in the model (i.e. using a randomisation algorithm that assumes uniform probability of species occurrence across sites). Note that for these analyses there will still be variability in species richness between sites, since species occupy sites randomly, but this variability will not necessarily reflect the variability in the observed data. This was done using 289 presence-absence datasets available in the nestedness temperature program (Atmar & Patterson 1993). Two datasets were not analysed from the original collection of 291, since for these it was not possible to randomise species co-occurrences. Note that many of these datasets relate to distributions of species across islands, although some are from continuous areas of habitat. The C-score was used to assess the degree of structure in these matrices. This metric is a standardised measure of the number of checkerboard units present in a matrix, i.e. the number of pairs of sites and pairs of species for which each species occurs only once and at a different site (Stone & Roberts 1990). If the C-score is higher than expected by chance, then pairs of species tend to be segregated across sites. The C-score was calculated for each matrix *(nestedchecker* function in the R package *vegan;* Oksanen et al. 2018). Each matrix was then randomised 5000 times while maintaining occurrences per species, but assigning species occurrences to different sites with equal probability. This was done with algorithm *c0* in the R *oecosimu* function (Oksanen et al. 2018) and results in matrices with identical species occurrence distributions and identical matrix fill to the observed dataset, but with each individual species occurrence being assigned to a site at random (unless that site is already occupied). P-values and standardised effect sizes were calculated on the basis of these null distributions. For each matrix I also calculated the mean absolute deviation (MAD) in species richness between sites as an easily interpretable measure of variability in species richness. The species richness MAD was then used to predict C-score standardised effect sizes (SES values) using a linear model. In the model I also included the matrix fill, the number of sites, and the number of species, since these are known to affect probability of pattern detection (Fayle & Manica 2010; Pitta et al. 2012; Ulrich & Gotelli 2007), and MAD is likely to be greater in larger matrices, all other things being equal. To test for any nonlinearity in the relationships between predictors, which might make them unsuitable for use in this kind of model, I plotted MAD against number of sites and number of species, and failed to find any nonlinear patterns (Supplementary Figure 1). To achieve normally distributed residuals in this analysis and in all following analyses, the response (C-score SES) and all predictors with the exception of matrix fill were log transformed.

I carried out a second set of analyses to assess whether detection of non-random patterns of species co-occurrence could arise solely from high variability in species richness with species otherwise co-occurring at random. To do this I generated a series of 289 matrices for which the row and column sums (and therefore also matrix fill) were fixed to those from the real 289 matrices, but in which species occurrences were randomised. This meant that for each matrix the distribution of species richness between sites, occurrences between species, matrix fill, and matrix size were all identical to the corresponding real matrix. This was done by running the *quasiswap* algorithm in the R package *vegan* (Oksanen et al. 2018) on the corresponding real dataset, and then conducting null model analyses in the same way as described in the preceding paragraph. I also ran all of the analyses presented here using the *curveball* randomisation algorithm in *vegan* (Strona et al. 2014), to check that these two commonly used fixed-fixed algorithm give similar results. The results of these analyses are almost identical to those using the *quasiswap* algorithm and hence only the latter are presented in the main text (see Supplementary Figures 2 and 3 for plots using *curveball),* I then used the same linear modelling framework as described above for the real datasets to assess the effects of species richness MAD, matrix fill, the number of sites, and the number of species on C-score standardised effect sizes, in the absence of any interactions between species. Null model tests of these pre-randomised matrices should give significant results (i.e. result in type 1 errors, since species co-occurrence is known to be random) at a rate equal to the critical p-value. In order to test these type 1 error rates, the total number of matrices (out of 289) giving significantly non-random results was tested against a binomial distribution with P(significant)=0.1 (note that *oecosimu* uses p=0.1 as a critical value for two-tailed tests).

### What is the effect of including variability in species richness between sites within a dataset in null models on detection of co-occurrence patterns?

Including observed variability in species richness in null models should decrease the probability of detecting non-random species co-occurrence. This is because, first, for communities in which differences between species have generated non-uniform patterns of species richness, the power to detect this effect will be decreased (the Narcissus effect, see Introduction). Second, in communities in which species richness deviates from the distribution expected with equal probabilities of species occupation across sites as a result of processes unrelated to species co-occurrence, for which an equiprobable model therefore incorrectly detected non-random co-occurrence patterns, these errors will be corrected. I tested the degree to which accounting for differences in species richness between sites in null models altered the outcomes of analyses of real and simulated communities. The null models of species co-occurrence and linear models linking MAD and standardised effect sizes were rerun on the real and pre-randomised datasets, but null matrices for were generated using a fixed-fixed randomisation algorithm that does not allow species richness in the null matrices to deviate from that in the observed matrix (algorithm *quasiswap* in R *oecosimu* function, see Supplementary Figure 3 for parallel analysis using the *curveball* algorithm). Note that the pre-randomised datasets were the same as those used in the first section, being generated using the *quasiswap* algorithm that constrains both row and column sums. Hence the second of these analyses involving matrices pre-randomised using the *quasiswap* algorithm, and then analysed using the *quasiswap* algorithm as a null model are expected to give significant results no more than would be expected at random, and are presented here for completeness only. Type 1 error rates were tested using binomial tests, as described above.

## Results

### Does variability in species richness between sites within a dataset correlate with detection of deterministic species co-occurrence patterns?

Of the 289 real matrices tested, 218 (75%) showed significant deviations in the C-score from null models for which observed richness variability between sites was not included (Table 1). The C-score standardised effect size (SES) was more negative with increasing variation in species richness between sites (Linear model: mean absolute deviation: t_4,284_=-3.54, P<0.001, Figure 1A,B). That is, detecting non-random species co-occurrence was more likely if there was high variation in species richness between sites within a dataset (Figure 1A). This was still the case after taking into account negative relationships between SES and number of species, number of sites, and matrix fill (species: t_4,284_=-28.75, P<0.001; sites: t_4,284_=-3.03, P=0.003, fill: t_4,284_=-9.50, P<0.001). The majority of observed matrices that deviated significantly from null models had fewer checkerboard units than would be expected at random (205 of 218 non-random results; 94%), meaning that there were low levels of segregation between species.

**Figure 1.**
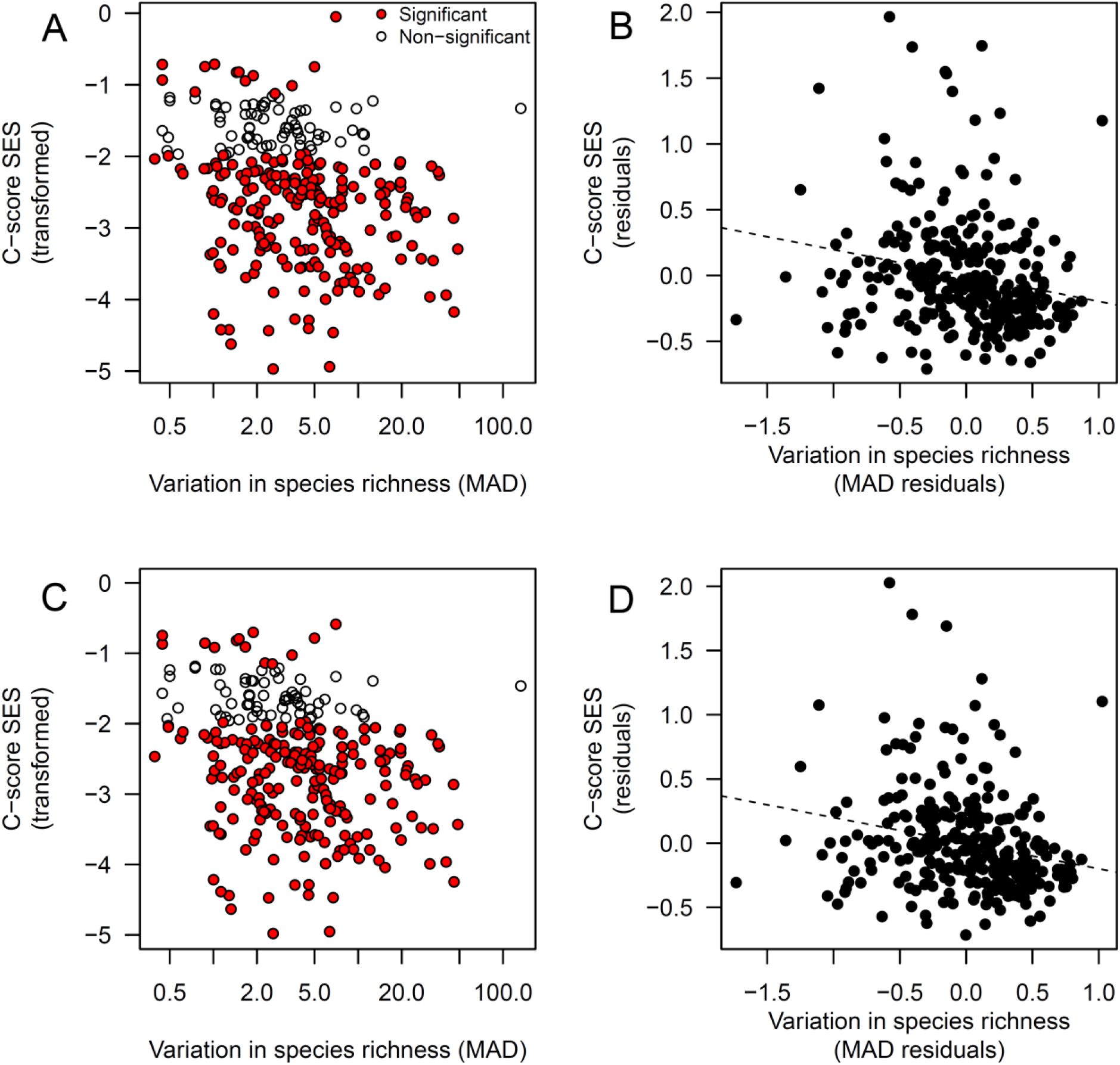
Relationship between variability in species richness and deviation from random species assembly when observed variation in species richness is not accounted for in the null model. (A) With increasing variability in species richness between locations within a site (Mean Absolute Deviation, MAD), the detected strength of deviation from random species assembly in real matrices increases when this variation in species richness is not included in the model (fixed-equiprobable randomisation algorithm). Red points denote statistically significant deviations from random species assembly. (B) This is still the case when the effects of number of sites, number of species and matrix fill are taken into account (partial regression plot). (C) When the species richness and number of sites are fixed to those from the 289 real matrices, but with species being placed at random to each other to create datasets, the same negative relationship between MAD and SES is observed. (D) Partial regression plot displaying this pattern.

**Table 1.**
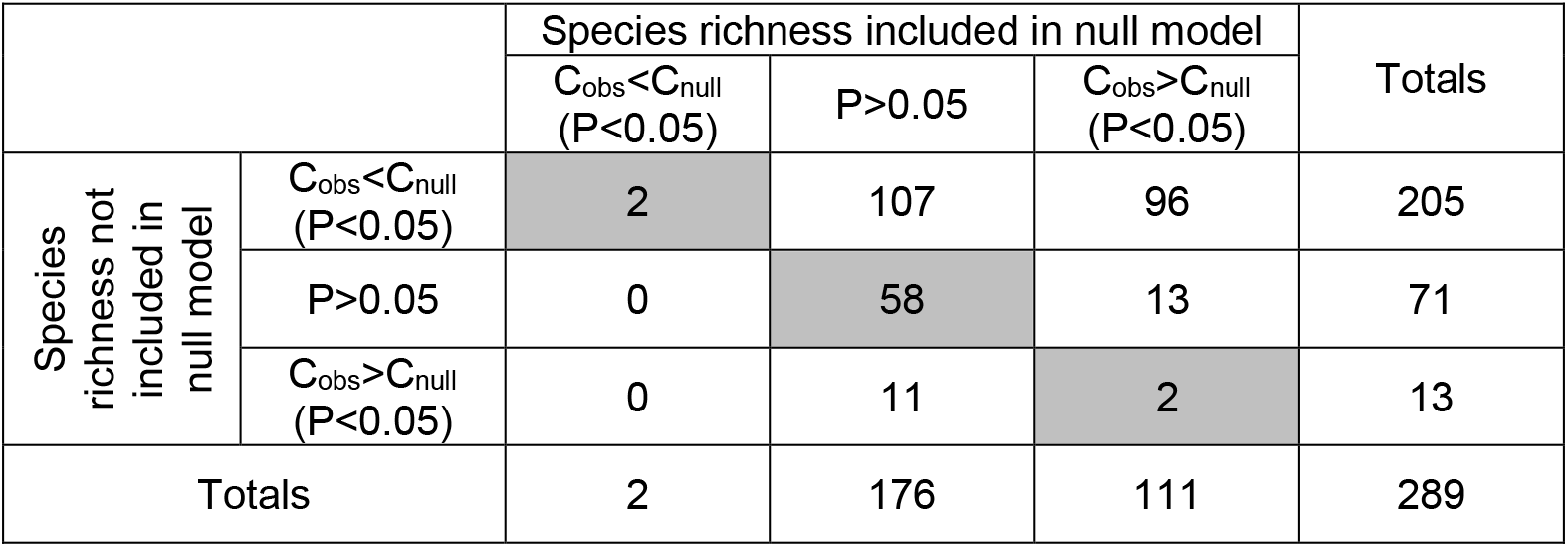
Influence of restricting randomisation algorithm to observed species richness across sites on significance and direction of results from null models of co-occurrence. The effects of including species richness in a null model of species co-occurrence (fixed-fixed randomisation algorithm) or not (fixed-equiprobable algorithm) on frequencies of significant test results for real datasets. Where the observed C-score is significantly higher than the null distribution of C-scores (C_obs_>C_null_), then there is more segregation between species than expected at random, and vice versa for C_obs_<C_null_.

For randomly generated matrices with identical species richness and species occurrence distributions to the real matrices, 223 of the 289 sites (77%) showed non-random species co-occurrence (one tailed binomial test against P(significant)=0.1: P<0.001). As was the case with the real matrices, the C-score standardised effect size was more negative with increasing variation in species richness between sites (Linear model: mean absolute deviation: t_4,284_=-3.94, P<0.001, Figure 1C,D). This was still the case after taking into account negative relationships between SES and number of species, number of sites, and matrix fill (species: t_4,284_=-31.69, P<0.001; sites: t_4,284_=- 3.43, P=0.001, fill: t_4,284_=-9.75, P<0.001). The high proportion of significant results indicates that analyses of these datasets, in which species co-occurred at random, suffered from extremely high levels of type 1 errors. If non-random co-occurrence between species had generated variation in species richness, then this analysis of randomised datasets and the first set of analyses should have given different results. This is because the signature of non-random co-occurrence that generated differences in species richness should still be present in the first set of analyses, but would have been removed by pre-randomisation for the second set of analyses. However, this was not the case, and results of the two sets of analyses were similar (205 real matrices vs. 210 pre-randomised matrices with C-score lower than expected at random, 13 real matrices vs. 13 pre-randomised matrices with C-score higher than random).

### What is the effect of including variability in species richness between sites within a dataset in null models on detection of co-occurrence patterns?

When using a randomisation algorithm that preserves the number of species per site in addition to the number of occurrences per species (fixed-fixed), for real matrices, 111 of the 289 sites (38%) showed non-random species co-occurrence. There was no longer a relationship between C-score SES and variation in species richness (Linear model: t_4,284_=0.17, P=0.866, Figure 2A,B). There were still positive relationships between C-score SES and number of species, number of sites, and matrix fill (species: t_4,284_=-7.70, P<0.001; sites: t_4,284_=2.86, P=0.005, fill: t_4,284_=2.64, P=0.009). The majority of datasets for which there was significantly non-random co-occurrence had a higher C-score than would be expected (111 of 113 non-random results, 98%), in contrast with the results for the fixed-equiprobable algorithm (not accounting for observed variation in species richness), which gave most matrices as having significantly lower C-score than would be expected (205 of 218 non-random results, 94%; Figure 1A, Figure 2A, Table 1).

**Figure 2.**
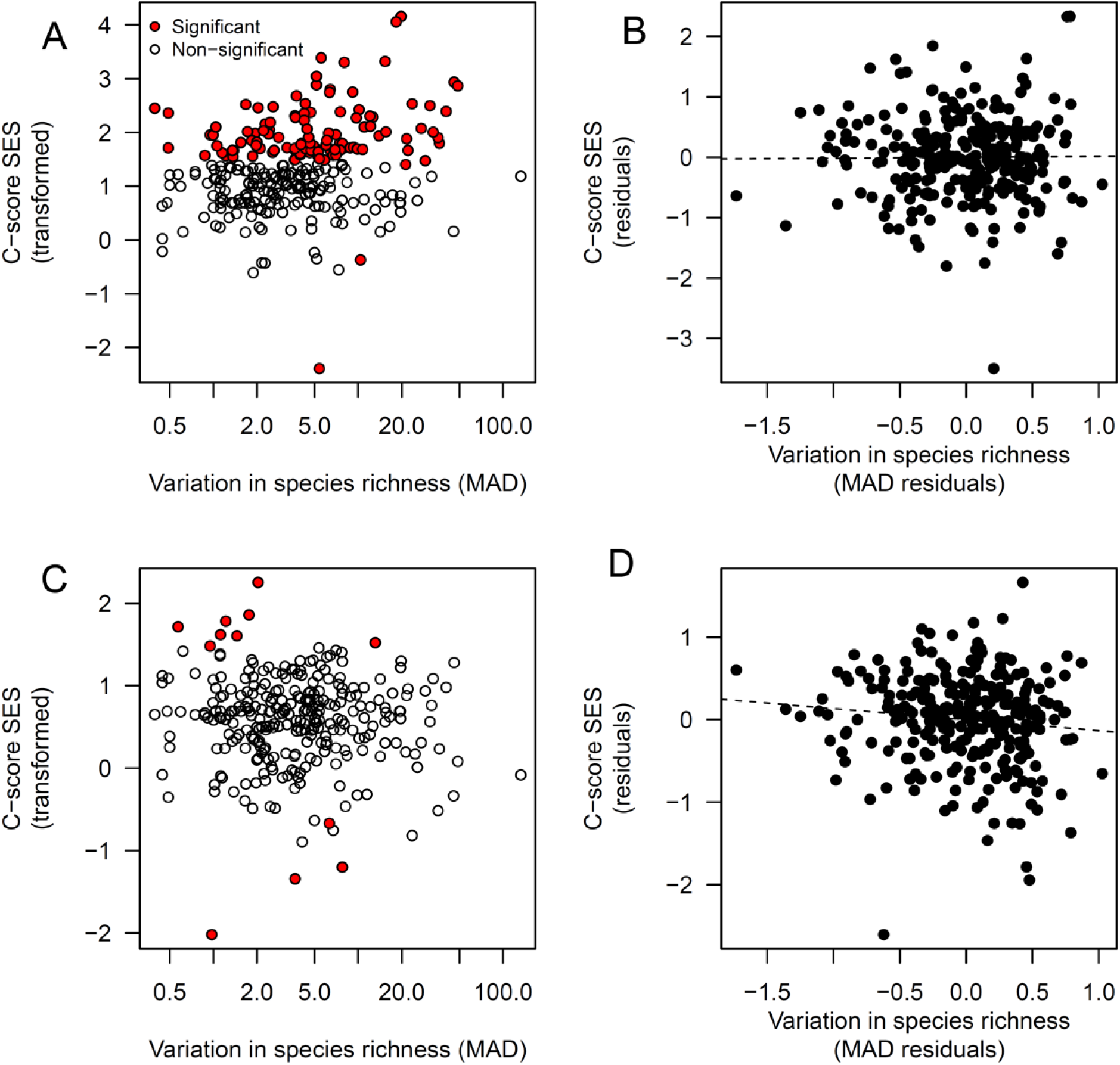
Relationship between variability in species richness and deviation from random species assembly when observed variation in species richness is accounted for in the null model. (A) When variation in species richness is included explicitly in the model of species co-occurrence (fixed-fixed randomisation algorithm) then for real matrices there is no relationship between variability in species richness between locations and the detected strength of deviation from random species assembly. (B) There is also no relationship when the effects of number of sites, number of species and matrix fill are taken into account (partial regression plot). (C) This is also true for the datasets for which the species richness and number of sites were fixed to those from the 289 observed matrices, but with species being placed at random to each other. (D) Partial regression plot displaying this pattern. Note that p-values are calculated from the raw null distribution, while standardised effect sizes are calculated from the variance in the null distribution. This means that occasionally some data points with more extreme SES values may be non-significant, while those with less extreme values are significant (e.g. in panel A).

As expected, for randomly generated matrices with identical species richness and species occurrence distributions to the real matrices, fixing the number of species per site in null models resulted in the loss of the relationship between C-score standardised effect size and variation in species richness between sites, although the relationship was only marginally non-significant (Linear model: mean absolute deviation: t_4,284_=-1.87, P=0.062, Figure 2C,D; compare to Figure 1C,D). Furthermore, there were also no relationships between SES and number of species, number of sites, and matrix fill (species: t_4,284_=-1.02, P=0.309; sites: t_4,284_=-0.09, P=0.173, fill: t_4,284_=-0.06, P=0.956). From the 289 sites, 12 (4%) showed non-random species co-occurrence (one tailed binomial test against P(significant)=0.1: P>0.999), indicating that type 1 error rates were within acceptable limits (10%) for these analyses. These results were expected because the same algorithm was used to generate the “observed” datasets and to create corresponding null models.

## Discussion

My results show that for this set of 289 real matrices, if null models are run without including observed variation in species richness, the majority of tests are significant, with most matrices having a smaller C-score (fewer checkerboard units) than would be expected at random. Furthermore, the greater the variation in species richness across sites, the larger the effect size. Such a pattern might be caused in two different ways. First, matrices with high variation in species richness might be those for which particular pairs of species co-occur non-randomly. In this case, since C-scores were lower than expected, this could be interpreted as evidence for ecological mechanisms that might lead to aggregation, such as mutualism, facilitation, or shared environmental preferences. It is easy to see how such mechanisms might give rise to sites with variable species richness. For example, if one “aggregator” species was present, which made the co-occurrence of many other species more likely, then this could generate sites with higher species richness. Conversely, if a strongly competitive species were present, which made co-occurrence with other species less likely, then this could generate sites with lower species richness. However, there is a second possible explanation for the tendency of matrices with high variance in richness to give significant results.

This alternative explanation is that variation in species richness is not related to non-random co-occurrence between species, and hence this variation has been incorrectly excluded from the null model, giving rise to false positive results (Type 1 errors). I explored this possibility by rerunning the same null model analyses, but with matrices pre-randomised, while maintaining species richness per site, and occurrences per species. These tests gave similar results to those conducted on the real datasets. Of 218 real datasets that tested significant using the fixed-equiprobable randomisation algorithm, 188 of these (86%) remained significant even when all structure other than species richness differences had been removed through a preliminary fixed-fixed randomisation (Figures 1a vs Figure 1c). It is therefore unclear whether the significant results for real datasets are driven entirely by variation in species richness between sites (unrelated to co-occurrence patterns between particular pairs of species), or if species really do co-occur non-randomly, and in the process of doing so, create variation in species richness. If the latter were the case, then one would expect that for matrices showing significantly low C-scores in the unconstrained analysis (e.g. where some hypothetical multi species mutualist had caused aggregations of species at high richness sites), then these significantly low C-scores should still be detectable even when richness distributions are fixed. I did not find this to be the case, since of the 205 matrices found to have a lower C-score than expected at random in the unconstrained analysis, only 2 (1%) also gave the same result in the fixed-fixed analysis. Furthermore, the results of the second set of tests are cause for concern, since they show that it is possible to incorrectly detect non-random co-occurrence patterns in random datasets if there is variation in richness between sites that is not accounted for in the null model. Hence for any particular null modelling exercise, if species richness is not included in the model, any significant results might be due either to non-random co-occurrence between species (the usual interpretation) or to variation in species richness that has not been included in the model. The fact that my results were similar between real and pre-randomised datasets indicates that the latter explanation is more likely to be correct in this case.

Why does variation in species richness (in the absence of non-random species co-occurrence) give rise to significantly different C-scores in the observed datasets from those predicted by the null model? I suggest that this relates to the way that variation in species richness between sites constrains the number of possible checkerboard units (Figure 3). For a given matrix, if variation in species richness is high, there are few possibilities for the formation of checkerboard units. However, species richness resulting from equal probability of site occupation (in this case represented as an extreme situation in which species richness is uniform, although note that if site occupancy is equiprobable, this would still lead to a case with variation in richness between sites) results in larger numbers of checkerboard units (on average). This holds for both structured and randomised matrices. Hence if a real matrix with high variation in species richness is randomised to create a null model under the assumption that species richness has lower variation, then the null model will have more checkerboard units than the observed dataset, independently of any non-random co-occurrence between species. In support of this speculation, an increase in C-scores for the real matrices used in this study was also observed when randomised using the c0 algorithm, but not when using fixed-fixed algorithms (Figure 4). Consequently, the real dataset will be interpreted as having a lower C-score than would be expected at random. Such results have been observed previously (Table 2 in Ulrich & Gotelli 2007), although have not been interpreted specifically in terms of the effects of variation in species richness between sites within a dataset. This potentially explains the large proportion of matrices for which the C-score was significantly lower (i.e. fewer checkerboard units) than would be expected under random community assembly in my analyses (Figure 1 A,C). Therefore, finding a significant result for a null model that does not explicitly account for variation in species richness could mean either that species co-occur non-randomly with respect to each other, or simply that species richness is more variable between sites than has been accounted for in the model (note that these explanations are not mutually exclusive).

**Figure 3.**
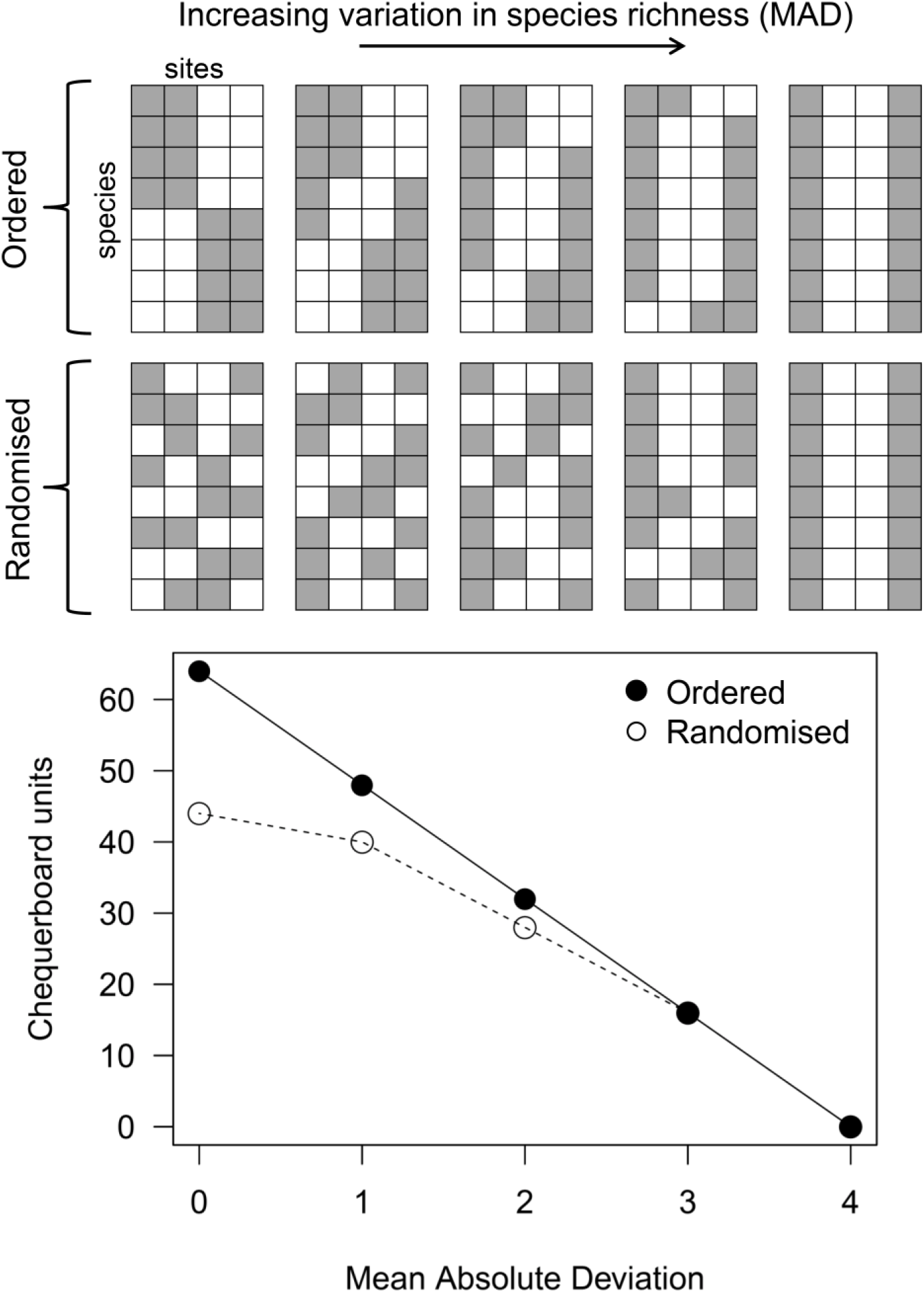
A demonstration for some small hypothetical datasets of the impact of variation in species richness on the possible number of checkerboard units. Assume that we have eight species distributed across four sites, and that each species always occurs at only two sites. Variation in site species richness can vary to give a range of values of mean absolute deviation (MAD), from zero to four. For both completely segregated matrices and randomised matrices the number of checkerboard units decreased with increasing variation in species richness. Note that median numbers of checkerboard units for 10,000 replications are plotted for randomised matrices, while only single example randomised matrices are plotted above.

**Figure 4.**
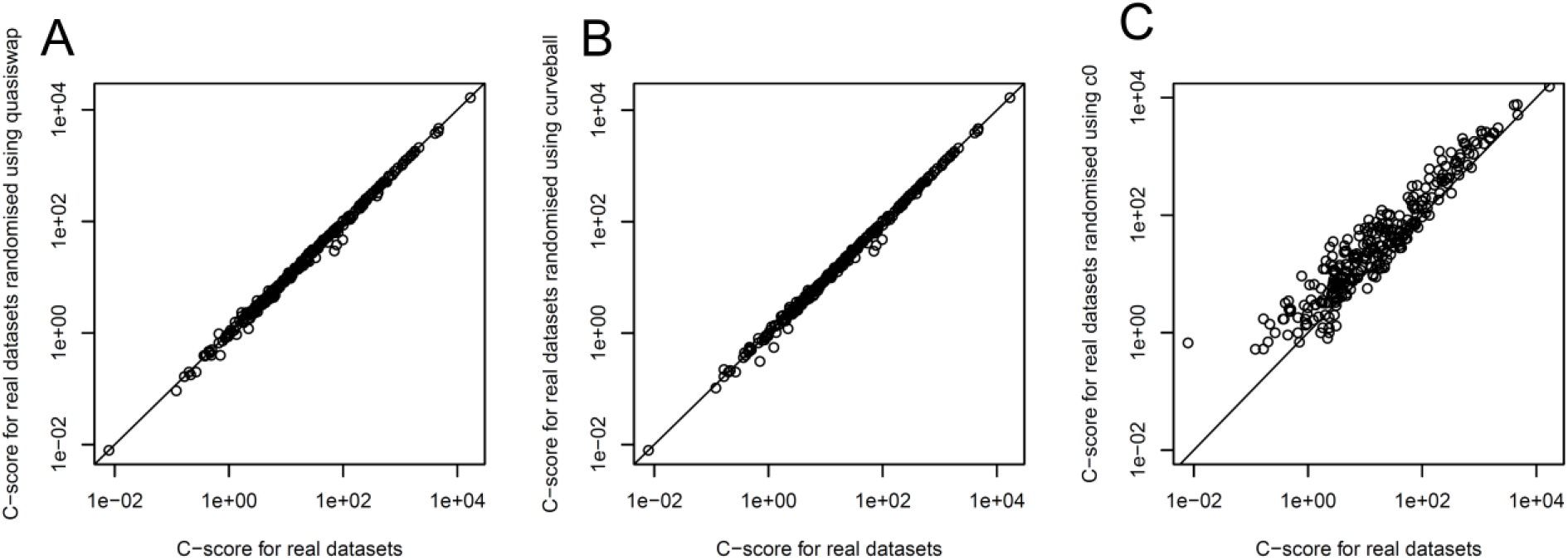
Impacts of randomisation on C-scores for real datasets for three algorithms. Relationship between C-scores for real datasets and single randomisations carried out using the two fixed-fixed algorithms quasiswap (A) and curveball (B), and the fixed-equiprobable algorithm c0 (C). Lines denote the scenario in which randomisation does not affect the C-score. Note the large proportion of points above the line when datasets are randomised using the c0 algorithm.

In order to avoid this issue, one approach is to constrain the null model species richnesses to those from the observed dataset. This allows detection of non-random co-occurrence within the pre-existing species richness distribution. Using this approach, I found that standardised effect size no longer correlated with variance in species richness, the proportion of significant results was lower, and the majority of those that were significant indicated that the observed dataset had a higher C-score than would be expected at random. Worryingly, of the 289 real datasets, 96 (33%) showed significant but *opposing,* patterns of species co-occurrence when the two different randomisation algorithms were used. For all of these, the C-score was lower than expected when using the fixed-equiprobable algorithm, and higher than expected when using the fixed-fixed algorithm. This indicates that some patterns of segregation between species were previously masked by not accounting for variation in species richness, which instead resulted in reporting of patterns of species aggregation. Effectively, assuming that species occurrence probability does not differ between sites in a null model results in real high richness sites being interpreted as species aggregations, while constraining null model species richness to that observed allows for the possibility that within a varying richness distribution, there might be some pairs of species that are segregated (or aggregated). This mechanism potentially explains previous results in which the direction of the reported species co-occurrence pattern depended on the randomisation algorithm used (Hétérier et al. 2008; Rooney 2008). My result also emphasizes the importance of specifying the null hypothesis under consideration (and hence choice of algorithm use), since the two most commonly used choices (fixed-equiprobable and fixed-fixed) give strong effects in the opposite direction for one in three datasets.

Taken together, these results are a cause for concern, because they suggest that the outcome of null modelling of species co-occurrence can depend greatly on the randomisation algorithm used. The same data could hence be interpreted as indicating entirely different patterns, for example species segregation as a result of competition, rather than species aggregation as a result of facilitation. Matrices that show high variation in species richness between sites are always likely to give significant results when tested against a null model that assumes low variance in richness, regardless of the way that pairs of species co-occur. This kind of bias may also explain the high type 1 error rates of algorithms that do not constrain species occurrences (73-76%; Gotelli 2000), since observed matrices may have high variation in species occurrences (i.e. they have skewed relative occurrence distributions). Although I only tested these effects with the C-score metric, it is likely that these differences in richness variation between observed and simulated matrices will affect the calculation of other metrics of co-occurrence. Researchers need to think carefully about how the value of their co-occurrence metric of choice is likely to change when matrices are randomised.

However, the possibility that variation in species richness might be generated by species interactions should also not be discounted, and understanding the links between pairwise species interactions and larger scale community patterns is likely to a fruitful avenue of future research. Indeed, an alternative approach that avoids the problems described here is to construct a series of hypothesised mechanistic models (as well as a null model) and use the results of these models as support for presence or absence of particular species assembly rules (Fayle et al. 2015).

In line with previous recommendations (Gotelli 2000; Ulrich & Gotelli 2007) researchers should have a strong rationale for using any randomisation algorithm other than one that fixes both the species richness distribution, and the number of occurrences per species. The analyses here demonstrate that use of multiple randomisation algorithms on the same datasets can be useful, if outputs are interpreted carefully. More broadly, these results illustrate how the comparison of a single alternative hypothesis to a null model is not a strong analytical strategy, since multiple mechanisms might cause deviations from the null model, including those not considered as part of the alternative hypothesis. A more powerful approach is the postulation of multiple competing alternative hypotheses, and the construction of mechanistic models of species co-occurrence based on these hypotheses to allow direct comparison of their explanatory power (Gotelli & Ulrich 2012).

## Acknowledgements

I am grateful to Vojtech Novotny and Kalsum Mohd Yusah for comments on this manuscript, to Andrea Manica for stimulating discussions about null modelling, and to the anonymous reviewers who have provided feedback on this manuscript. Funding was provided by a standard grant from the Czech Science Foundation (19-14620S).

## Data Accessibility

All datasets used in this paper were downloaded as part of the nestedness temperature programme (to which TMF is not affiliated), which is no longer maintained, but remains available in archived form (https://web.archive.org/web/20160819062742/https://web.archive.org/web/20160819062742/www.fieldmuseum.org/nestedness-temperature-calculator-program). All model outputs and code are available online at www.zenodo.org(https://doi.org/10.5281/zenodo.4244215).

