## Supplementary Figures 1-3 for "Varying richness need not imply non-random species co-occurrence: implications for specifying null models"

Short title: Species richness and co-occurrence

Author: Tom M. Fayle*

Address: Biology Centre of Czech Academy of Sciences, Institute of Entomology, Branišovská 31, 370 05 České Budějovice, Czech Republic; Institute for Tropical Biology and Conservation, Universiti Malaysia Sabah, Kota Kinabalu, Sabah, Malaysia

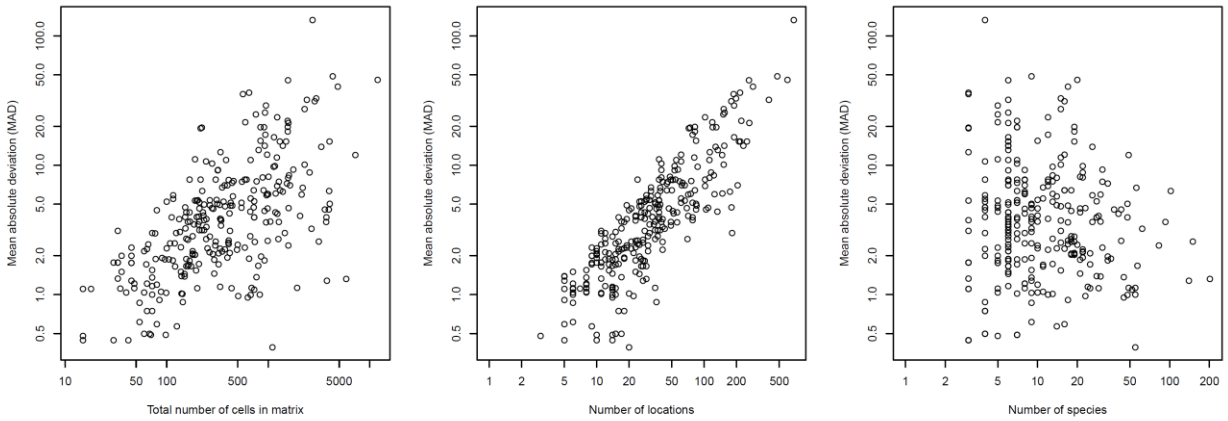


Supplementary Figure 1. Relationships between Mean Absolute Deviation and matrix size, number of locations, and number of species for all real datasets analysed.

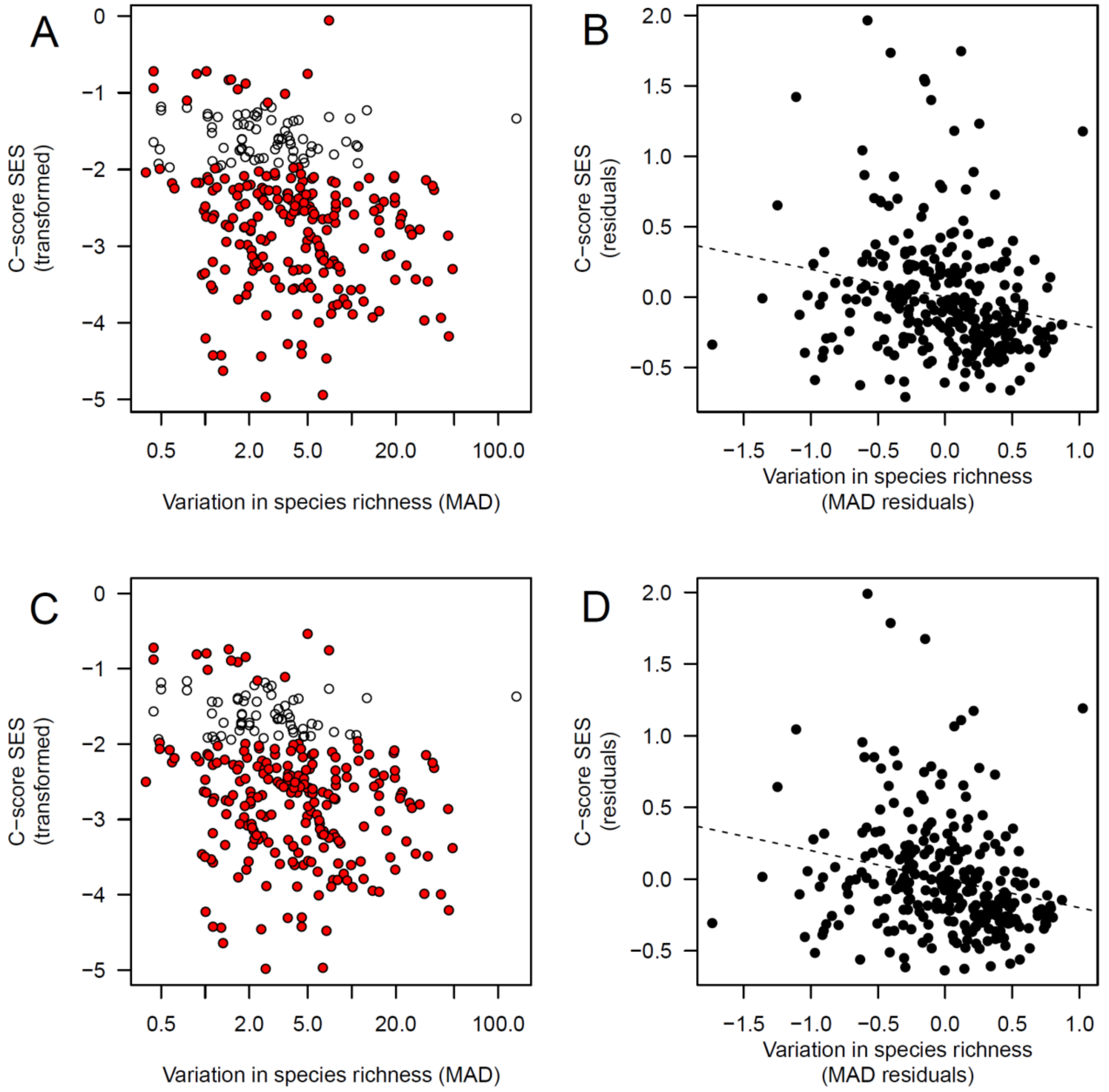


Supplementary Figure 2. Analyses presented in Figure 1 of the main text, but using the *curveball* randomisation algorithm rather than the *quasiswap* randomisation algorithm where applicable throughout.

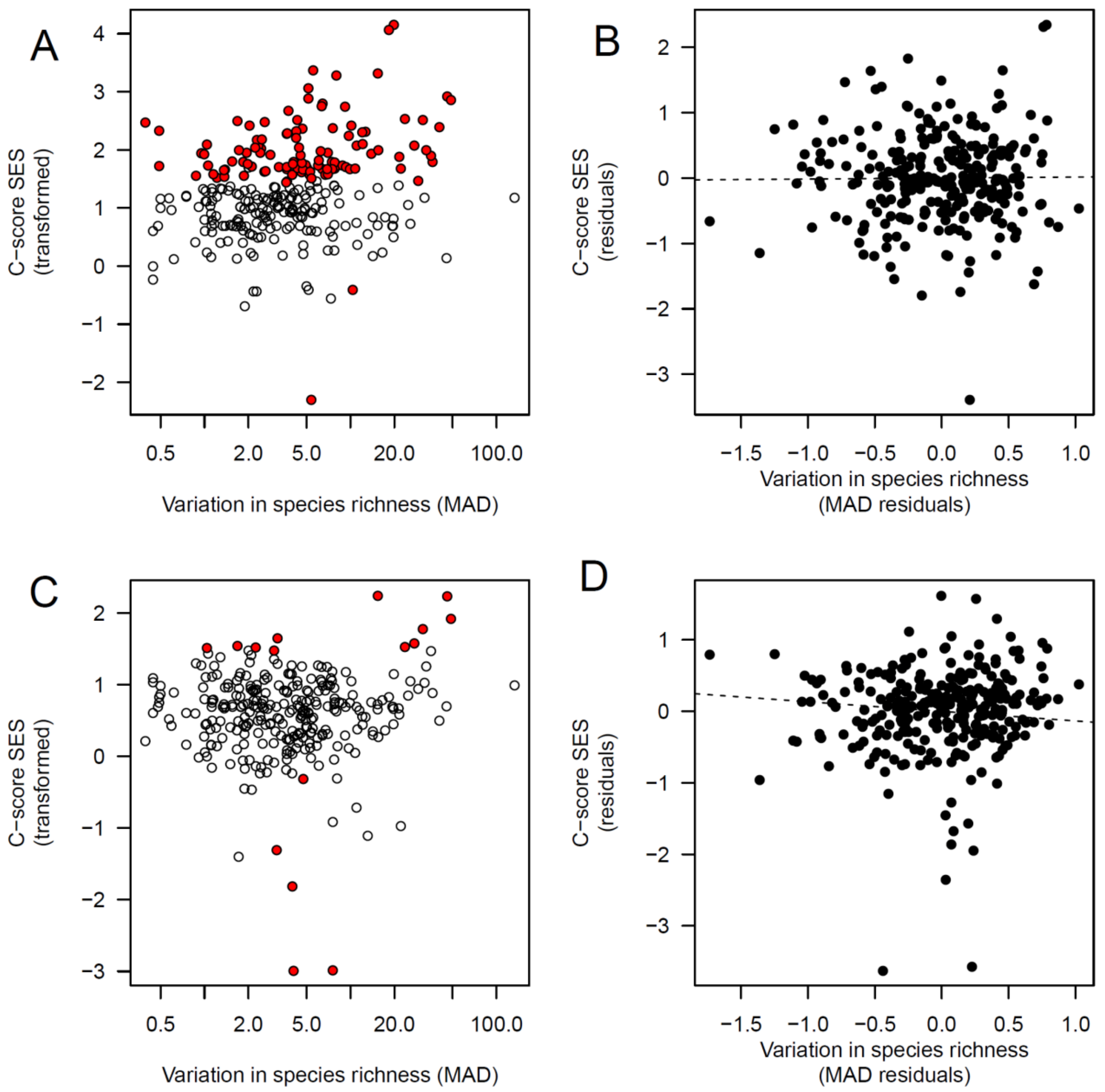


Supplementary Figure 3. Analyses presented in Figure 2 of the main text, but using the *curveball* randomisation algorithm rather than the *quasiswap* randomisation algorithm where applicable throughout.
